# Loss of the Reissner Fiber and increased URP neuropeptide signaling underlie scoliosis in a zebrafish ciliopathy mutant

**DOI:** 10.1101/2019.12.19.882258

**Authors:** Christine Vesque, Isabelle Anselme, Guillaume Pezeron, Yasmine Cantaut-Belarif, Alexis Eschstruth, Morgane Djebar, Diego López Santos, Hélène Le Ribeuz, Arnim Jenett, Hanane Khoury, Joëlle Véziers, Caroline Parmentier, Sylvie Schneider-Maunoury

## Abstract

Cilia-driven movements of the cerebrospinal fluid (CSF) are involved in zebrafish axis straightness, both in embryos and juveniles [1, 2]. In embryos, axis straightness requires cilia-dependent assembly of the Reissner fiber (RF), a SCO-spondin polymer running down the brain and spinal cord CSF-filled cavities [3]. Reduced expression levels of the *urp1* and *urp2* genes encoding neuropeptides of the Urotensin II family in CSF-contacting neurons (CSF-cNs) also underlie embryonic ventral curvature of several cilia motility mutants [4]. Moreover, mutants for *scospondin* and *uts2r3* (a Urotensin II peptide family receptor gene) develop scoliosis at juvenile stages [3, 4]. However, whether RF maintenance and URP signaling are perturbed in juvenile scoliotic ciliary mutants and how these perturbations are linked to scoliosis is unknown. Here we produced mutants in the zebrafish ortholog of the human *RPGRIP1L* ciliopathy gene encoding a transition zone protein [5–7]. *rpgrip1l*^-/-^ zebrafish had normal embryogenesis and developed 3D spine torsions in juveniles. Cilia lining the CNS cavities were normal in *rpgrip1l*^-/-^ embryos but sparse and malformed in juveniles and adults. Hindbrain ventricle dilations were present at scoliosis onset, suggesting defects in CSF flow. Immunostaining showed a secondary loss of RF correlating with juvenile scoliosis. Surprisingly, transcriptome analysis of *rgprip1l* mutants at scoliosis onset uncovered increased levels of *urp1* and *urp2* expression. Overexpressing *urp2* in *foxj1*-expressing cells triggered scoliosis in *rpgrip1l* heterozygotes. Thus, our results demonstrate that increased URP signaling drives scoliosis onset in a ciliopathy mutant. We propose that imbalanced levels of URP neuropeptides in CSF-cNs may be an initial trigger of scoliosis.

## RESULTS

### Rpgrip1l zebrafish mutants develop scoliosis and show cilia defects at juvenile stage

To study the mechanisms of scoliosis appearance upon cilia dysfunction we made use of a zebrafish deletion mutant in the *rpgrip1l* gene encoding a ciliary transition zone protein (mutant generation described in Supplementary Figure S1). *rpgrip1l*^Δ/Δ^ embryos were straight and did not display any additional defects found in many ciliary mutants such as kidney cysts, randomized left-right asymmetry or retinal anomalies [8–10] (Figure 1 and Supplementary Figure S1). *rpgrip1l*^Δ/Δ^ animals developed scoliosis during juvenile stages (3 weeks postfertilization (wpf) to 12 wpf), initiating by slight upward bending of the tail (tail-up phenotype) and progressing toward severe curvature (Figure 1A, B) with 90% penetrance in adults (100% in females and 80% in males) (Fig.1 C). Micro-computed tomography (μCT) at two different stages (5 wpf and 23 wpf) confirmed that spine curvature was tridimensional and initiated by a tail-up and showed no evidence of vertebral malformation or fracture (Figure 1 and Supplementary Movie 1). The *Ftm/rpgrip1l* null mouse mutant displays a severe embryonic phenotype with cilia loss in the nervous system and abnormal cilia in other structures [11,12]. In contrast, cilia appeared normal in zebrafish *rpgrip1l*^Δ/Δ^ embryos and larvae (Fig. 1G, H and Supplementary Figure S1), consistent with normal body axis geometry. On scanning electron microscopy (SEM) of the adult brain ventricles (Figure 1I-S), cilia of *rpgrip1l*^Δ/Δ^ multiciliated brain ependymal cells were sparse and disorganized (Figure 1I, L-N) compared to controls (Figure 1J, K). In the hindbrain, cilia of monociliated ependymal cells showed abnormal, irregular structures, with bulged and thinner parts (Fig. 1O-S). Cilia anomalies were already found in the spinal cord central canal (CC) at juvenile stages. Cilia were reduced in number and globally increased in length and showed strongly reduced Arl13b content (Figure 1 T-V’’ and Supplementary Figure 1L, M). Thus, *rpgrip1l*^Δ/Δ^ mutants show cilia defects appearing after embryogenesis and develop spine curvatures in juveniles.

**Figure 1:**
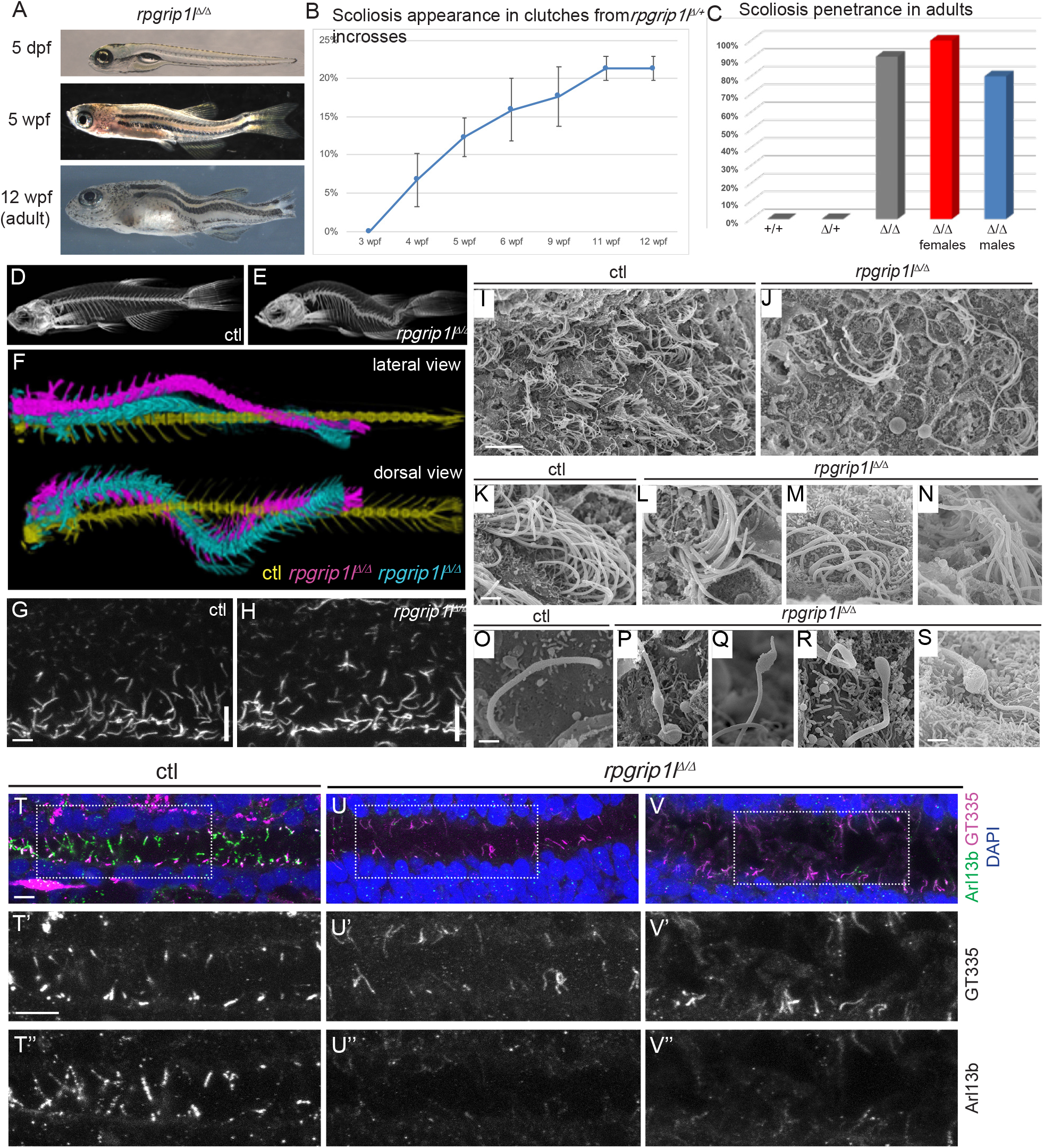
*rpgrip1l*^Δ/Δ^ zebrafish develop scoliosis and show cilia defects at juvenile stages. A) Representative *rpgrip1l*^Δ/Δ^ fish at 5 days post-fertilization (dpf) (4 mm body length, larvae), 5 wpf (1 cm body length, juveniles) and 12 wpf (2.5 cm body length, adults), showing absence of defects in embryos, onset of spine curvature (tail up) in juveniles and scoliosis in adults. B) Graph showing the dynamics of scoliosis appearance in *rpgrip1l*^Δ/+^ incrosses (total 4 clutches, 252 fish). C) Scoliosis penetrance in adults. D, E) Micro-computed tomography (μCT) scans of 4 control siblings (D) and 4 *rpgrip1l*^Δ/Δ^ (E) adult fish. F) Dorsal and lateral views of the superimposed spines of one control (yellow) and two *rpgrip1l*^Δ/Δ^ (pink, cyan) adult fish illustrating the 3D spine curvatures in mutants. G,H) Immunofluorescence for Acetylated tubulin in the neural tube of 2 dpf control (G) or *rpgrip1l*^Δ/Δ^ (H) embryos. White vertical lines indicate the floor plate I-S) Scanning electron microscopy of the brain ventricles of control (I, K, O) and *rpgrip1l*^Δ/Δ^ (J, L-N, P-S) fish showing ciliary defects in midbrain ependymal multiciliated cells (I-N) and in hindbrain monociliated cells (O-S). Scale bars: 3 μm for G,H, 10 μm for I,J; 1 μm for K-S and 10 μm for T-V’’.

### Rpgrip1l^Δ/Δ^ juveniles show ventricular dilations and loss of the Reissner fiber at scoliosis onset

Ciliary beating is an essential actor of CSF flow and of ventricular development in zebrafish larvae [13]. Thus, ciliary defects in the brain of *rpgrip1l*^Δ/Δ^ adults and juveniles could lead to abnormal ventricular volume and content. To determine whether ventricular dilations could be at the origin of scoliosis, we analyzed ventricular volume at the onset of spine curvature (Figure 2A-G). Ventricular reconstruction was performed on cleared brains of 4 control and 4 *rpgrip1l*^Δ/Δ^ (3 tail-up and 1 straight) fish at 4 wpf, stained with ZO1 to highlight the ventricular border and with Dil to outline brain shape, focusing on the posterior midbrain and hindbrain ventricles (Figure 2A-D). We identified a significant increase in ventricle volume in *rpgrip1l*^Δ/Δ^ fish compared to controls, restricted to ventral regions of the caudal midbrain (ROI4.4) and hindbrain (ROI6). The non-scoliotic mutant fish showed a milder dilation than the scoliotic ones (green circles in Figure 2F, G), as confirmed by sections through the hindbrain ventricle (Fig. 2E, G). In conclusion, ventricle dilations were present in the midbrain and hindbrain of *rpgrip1l*^Δ/Δ^ fish, and were more pronounced in fish at a more advanced stage of spine curvature, suggesting that these dilations could participate in scoliosis onset.

**Figure 2:**
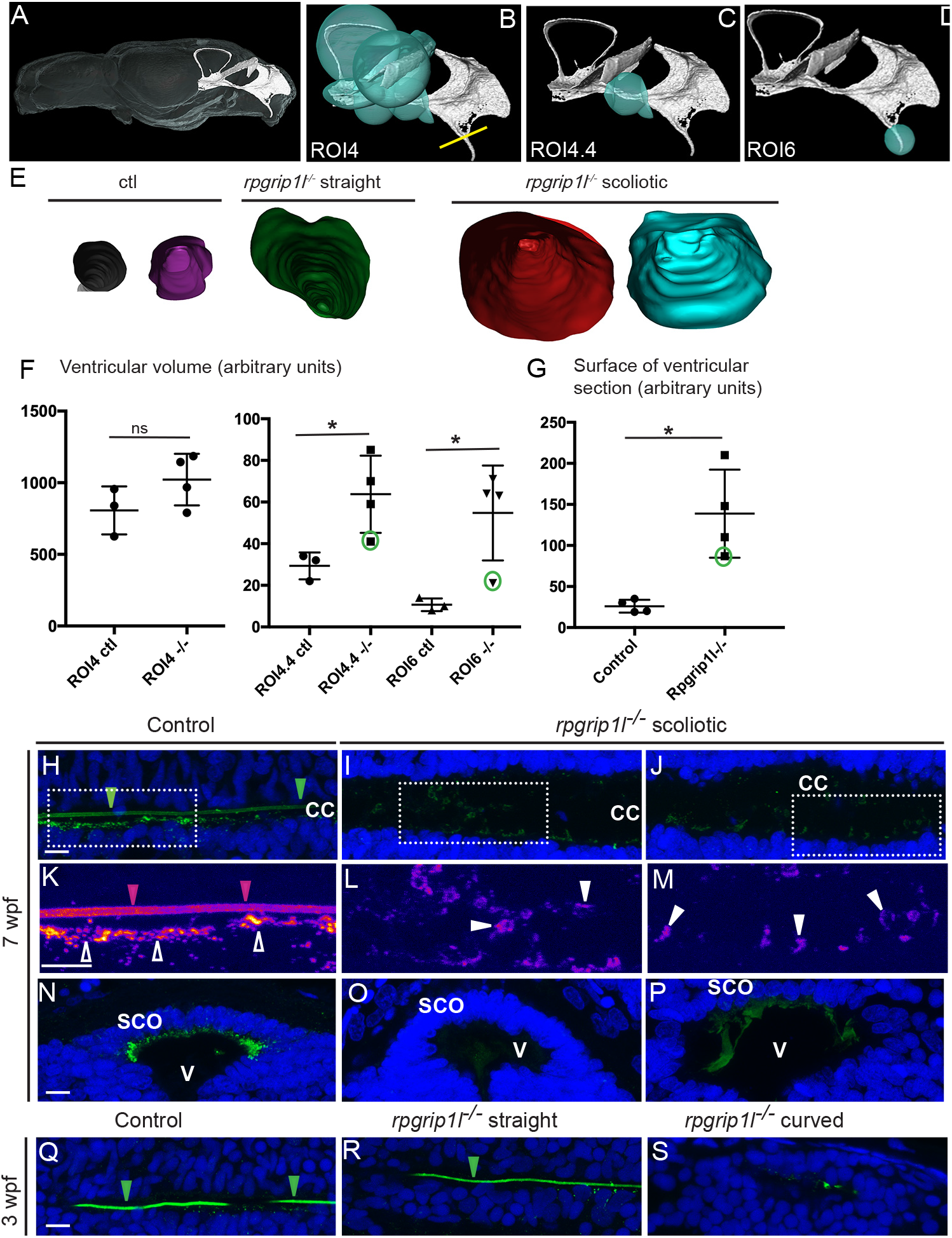
*rpgrip1l*^Δ/Δ^ juveniles show ventricular dilations and degeneration of the Reissner Fiber at scoliosis onset. A) Reconstruction of the posterior midbrain and hindbrain ventricles in a control transparized 5 wpf zebrafish brain stained with ZO1 antibody (ventricular surface) and DiI (global brain shape). B-D) Visualization of the ROIs shown in E. The yellow line in B indicates the level of optical sections in E. E) Optical transverse sections showing the caudal part of reconstructed ventricles of a control and two *rpgrip1l*^Δ/Δ^ fish, one straight and two tail-up. F) Graphs of the ventricle volume at the onset of scoliosis in 3 control and 4 *rpgrip1l*^Δ/Δ^ (one straight and three tail-up) fish. The dot circled in green corresponds to the straight mutant fish. G) Graph of the surface of the optical sections as illustrated in G. The dot circled in green corresponds to straight mutant fish. H-S) Visualization of the RF in sagittal sections of the trunk (H-M; Q-S) and of the brain at the level of the sub-commissural organ (SCO) (N-P) of juvenile fish, with a Sco-spondin antibody. H-P are 7 weeks post-fertilization (wpf) controls (H, K) or *rpgrip1l*^Δ/Δ^ (I, J, L, M, O, P) fish. Q-S are 3 wpf control (Q), *rpgrip1l*^Δ/Δ^ straight (R) or curved (S). K-M are higher magnifications of the regions boxed in H-J, respectively. Green arrowheads in H, Q and R and red arrowheads in K show the RF. Empty arrowheads in K show immunoreactive material in apical FP cells. At 7 weeks, the RF can be visualized in the central canal (CC) of controls (H, K). In the dilated CC of *rpgrip1l*^Δ/Δ^ scoliotic fish (I, J, L, M), the RF is not maintained but SCO-spondin-positive debris (white arrowheads in L, M) can be found in the CC. In the brain, SCO-spondin immunostaining is detected in controls at the apical side of SCO cells (N). This staining is strongly reduced in mutants, but large SCO-spondin debris are found in the third ventricle (V) (O,P). At three wpf, controls and straight *rpgrip1l*^Δ/Δ^ have a RF (Q, R) but the tail-up mutants have lost the RF (S). Scale bars: 10 μm in H-S.

The Reissner fiber (RF) is mainly composed of SCO-spondin secreted by the subcommissural organ (SCO) and floor plate (FP) (ref). To visualize it we used an antibody that labels the RF in zebrafish embryos [3,14]. The RF formed normally in *rgprip1l*^Δ/Δ^ embryos, as expected given the absence of embryonic curvature (Supplementary Figure 1). We could visualize the RF in wild-type adult brains in SEM (Supplementary Figure 2G-J). To study its maintenance in *rpgrip1l*^Δ/Δ^ juveniles, we immunostained spinal cord longitudinal sections. In 7 wpf control fish, the RF was visualized as a 1 μm diameter rod in the ventral part of the CC (Figure 2H, K, green arrowheads in Figure 2H). The antibody also stained the apical cytoplasm of FP cells (Figure 2H, K, empty arrowheads in Figure 2H) as well as the SCO in the caudal diencephalon (Figure 2N), showing that these two structures continue to produce and secrete SCO-spondin at juvenile stages. Thus, the RF is still produced and polymerized at juvenile stages, both in the brain ventricles and spinal cord CC, opening the possibility that growth and renewal of this structure beyond the larval stage could also depend on cilia integrity.

In 7 wpf *rpgrip1l*^Δ/Δ^ fish, the RF was absent and SCO-spondin-positive debris were abundant in the central canal (Figure 2I, J, L, M). We found a RF on 96% of the sections had a RF Transverse sections at the level of the diencephalon also revealed abnormally packed material in the ventricle (Figure 2O, P). To test whether the loss of RF could drive scoliosis, we observed this structure at 3 wpf in tail-up and straight *rpgrip1l*^Δ/Δ^ fish. All controls had a RF (found on 96% of the sections with a visible CC, 73 sections form 5 fish analyzed). All tail-up mutants had lost the RF (small remnants found on 5% of the sections, 40 sections with CC from 2 fish), while straight mutants had a RF (found on 98% of the sections, 45 sections with CC from 3 fish) (Figure 2Q-S). Thus, in *rpgrip1l*^Δ/Δ^ fish, the loss of RF correlated with scoliosis onset.

### Regulators of motile ciliogenesis, embryonic axis straightness and inflammation genes are upregulated in rpgrip1l^Δ/Δ^ juveniles

To get further insight into the mechanisms of spine curvatures in *rpgrip1l* mutants, we obtained the transcriptomes of the brain and trunk of 5 control (2 wt and 3 *rgprip1l*^Δ/+^) and 7 *rpgrip1l*^Δ/Δ^ juveniles (6 tail-up and 1 straight). Differential gene expression analysis was performed with one-way ANOVA. A complete annotated gene list is displayed in Supplementary Table 1. This analysis showed a large predominance of upregulated genes in the trunk (32 downregulated vs 1001 upregulated genes), and less in the brain (56 downregulated vs 235 upregulated genes) (Figure 3A). In the trunk, the most upregulated biological processes in GO term analysis included cilium movement, cilium assembly, cell adhesion, extracellular matrix organization and inflammation (Figure 3B). Many genes upregulated in the brain were also upregulated in the trunk, suggesting that, in the trunk, these genes were specific to the CNS (Figure 3C). Among the 15 most significant GO terms, 9 were related to cilium movement or cilium assembly (not shown). This was likely due to the 5-fold upregulation of *foxj1a*, encoding a master transcriptional activator of genes involved in ciliary motility [15,16] (Figure 3D). To confirm this hypothesis, we compared our dataset with those of targets of Foxj1 from [17–19]. 208 upregulated genes were targets of Foxj1, of whom 146 were direct targets (Supplementary Table 2). Another large series of genes overexpressed in mutants belonged to genes involved in inflammatory processes and immune response (Figure 3D), reminiscent of the *ptk7* scoliotic mutants [20].

**Figure 3:**
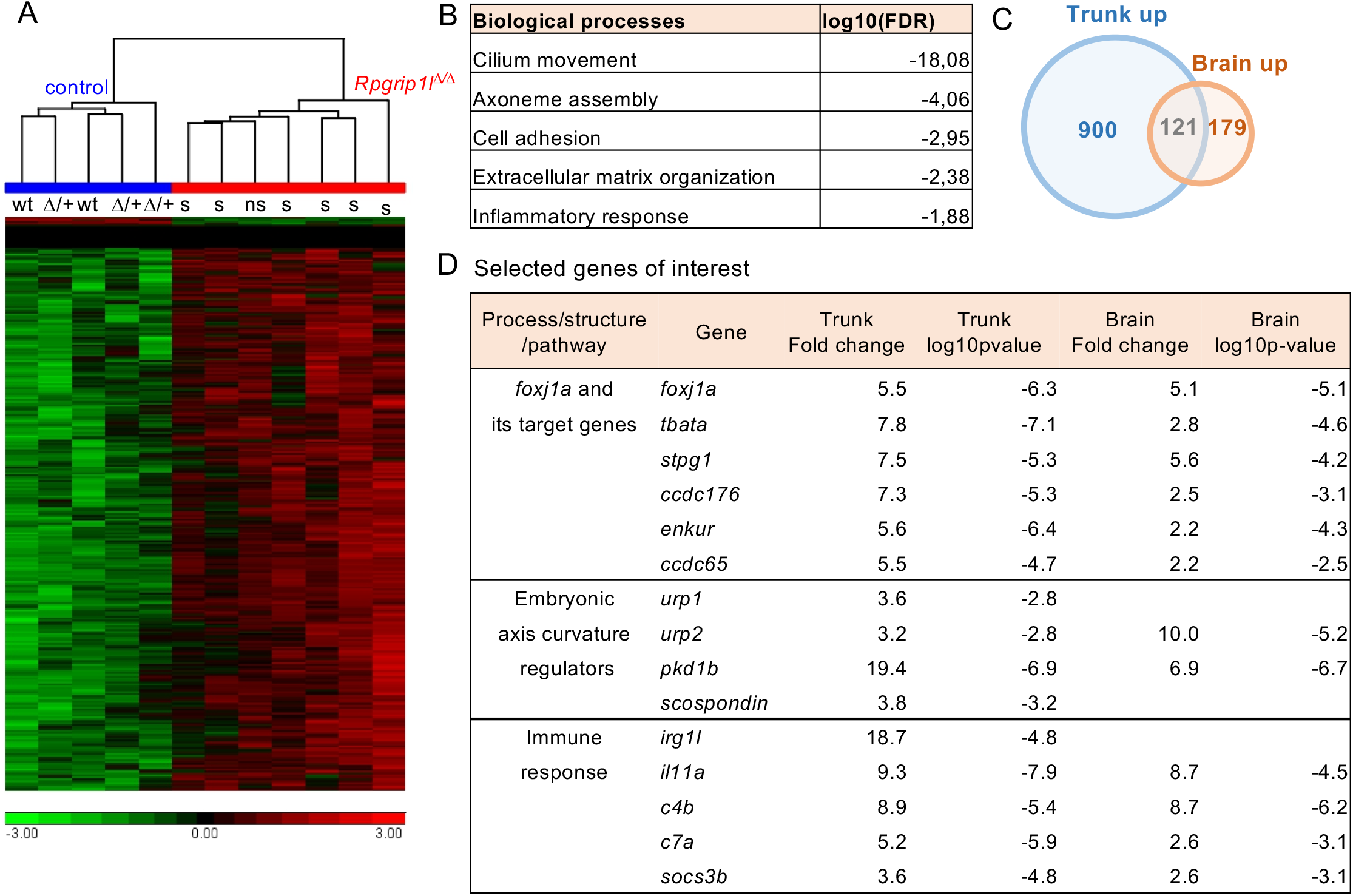
Comparative transcriptome analysis of the trunk and brain of control and *rpgrip1l*^Δ/Δ^ fish. A) Hierarchical clustering of the deregulated genes in the trunk (after removal of the skin and internal organs) of 2 wt, 3 *rpgrip1l^Δ/+^* and 7 *rpgrip1l*^Δ/Δ^ (6 tail-up and 1 straight) fish at 4 wpf (0.9 cm body length). B) Go term analysis of biological processes in the trunk samples. C) Vent diagram of the upregulated genes in brain and trunk samples, based on human Refseq, showing many common genes between the two tissues. D) Selected genes of interest upregulated in the trunk and/or brain of *rpgrip1*^Δ/Δ^ fish.

### Upregulation of URP signaling participates in scoliosis onset in *rpgrip1l* mutant fish

Several genes encoding regulators of embryonic axis curvature were also upregulated in *rpgrip1l*^Δ/Δ^ fish, among which *pkd1b* [21], *urp1* and *upr2* [4] and *scospondin* [3]. *urp1 and urp2* encode peptides of the Urotensin II family and are expressed in ventral spinal CSF-cNs [22,23]. Their downregulation underlies embryonic ventral curvature of several cilia motility mutants [4]. Thus, we wanted to confirm their overexpression and investigate its functional significance. qPCR analysis in 5 control and 5 *rpgrip1l*^Δ/Δ^ fish at the 3-4 wpf stage (0.8-0.9 cm) confirmed *urp2* overexpression in *rpgrip1l*^Δ/Δ^ fish and showed that this overexpression preceded scoliosis onset (Figure 4A). To investigate the tissue specificity of this up-regulation, we performed immunofluorescence with an antibody to a conserved domain of the Urotensin II peptide family [24,25]. In the zebrafish spinal cord, *urp1* and *urp2* are expressed in ventral CSF-cNs while other genes of the family are expressed in the caudal neurosecretory system [23,25,26]. Immunostaining on sagittal sections of control (B, C), straight (D, E) and scoliotic (F, G) *rpgrip1l*^Δ/Δ^ juvenile fish (3 wpf) showed that the proportion of URP-expressing cells was increased along the CC of straight *rpgrip1l*^Δ/Δ^ fish (Figure 4H) (this could not be analyzed in scoliotic fish because the sections rarely contained a complete portion of the CC). Moreover, the ratio of highly expressing CSF-cNs (full arrowheads) was significantly increased in scoliotic (but more variably in straight) *rpgrip1l*^Δ/Δ^ fish. To determine whether increased expression of URP1/2 peptides can trigger scoliosis, we injected a transgenic construct expressing *urp2* in *foxj1*+ cells, lining central nervous system cavities [27], in eggs from *rpgrip1l*^Δ/+^ crosses. Injected larvae displayed severe axis curvature (arrowhead) at 1-5 dpf (Figure 4J) and most of them died before 10 days. Among the survivor fish, 2/17 (12%) *rpgrip1l*^Δ/+^ heterozygotes became scoliotic whereas none of the non-injected *rpgrip1l*^Δ/+^ heterozygotes became scoliotic (Figure 4K). This indicates that overexpression of *urp2* in cells lining cavities of the central nervous system can trigger scoliosis onset in a sensitized background.

**Figure 4:**
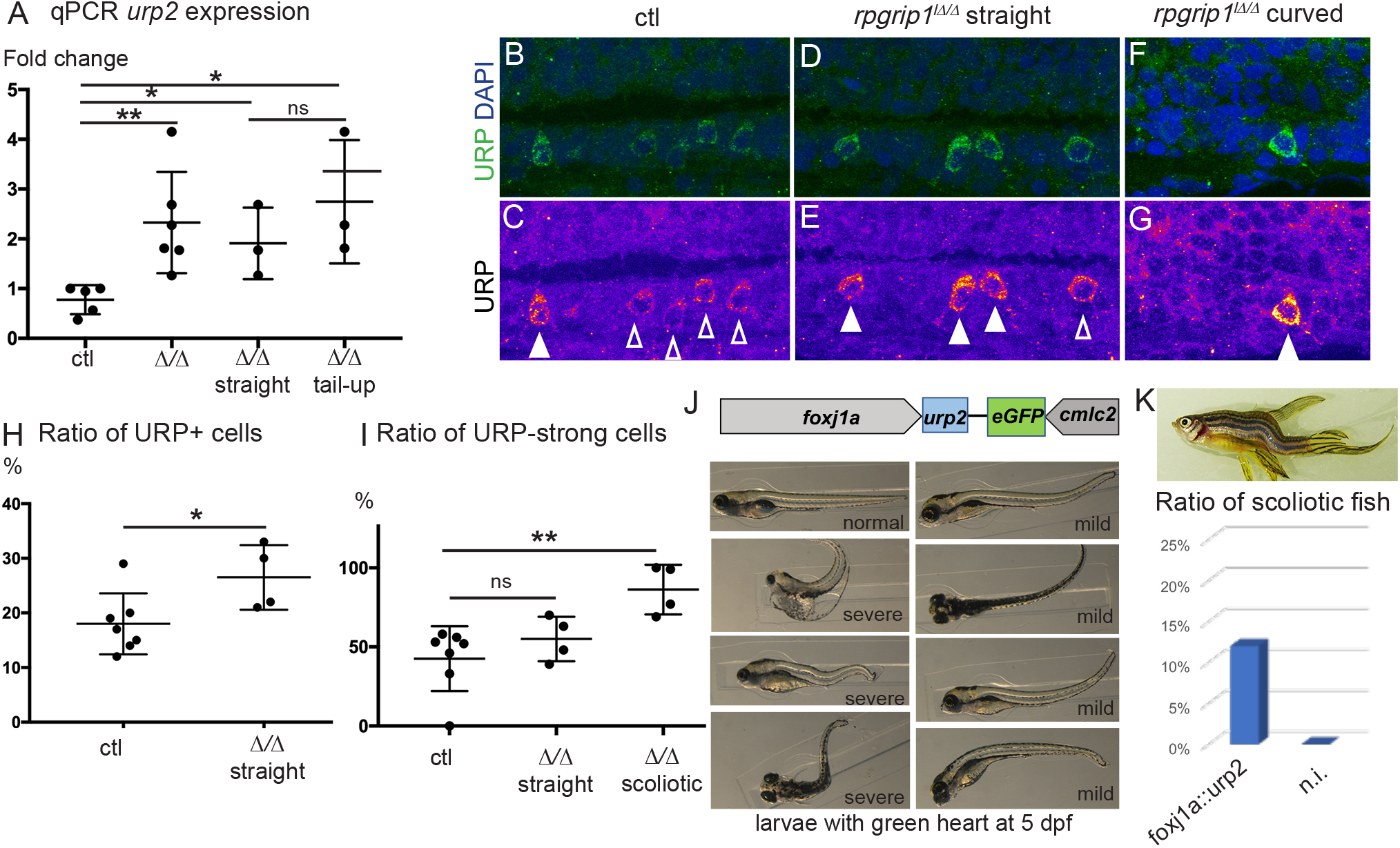
regulation of URP neuropeptide signaling underlies scoliosis onset in rpgrip1l mutants. A) qPCR confirms upregulation of *urp2* expression in *rpgrip1l*^Δ/Δ^ fish at the onset of scoliosis. Data are given in fold change compared to a control sibling chosen as standard. B-I) Immunostaining on sagittal sections of control (B, C), straight *rpgrip1l*^Δ/Δ^ (D, E) and curved *rpgrip1l*^Δ/Δ^ (F, G) juvenile fish (3 wpf) with an antibody to the Urotensin II/URP family of neuropeptides shows that the proportion of URP1/2-expressing CSF-cNs is increased along the central canal of straight fish (H), and the ratio of highly expressing cells (full arrowheads) is increased in scoliotic *rpgrip1l*^Δ/Δ^ fish (I). J) Injection of a transgenic construct expressing *urp2* in *foxj1+* cells in fertilized eggs from *rpgrip1l*^Δ/+^ crosses lead to axis curvatures in F0 injected fish. The construct also expressed eGFP in the heart (cmlc2 promoter) in order to assess for injection efficiency. Among injected larvae with green heart at 5 dpf, 36% (21/58) had a straight axis (labelled straight), 24% (14/56) had mild curvatures (mild, right images, the second from the top in a dorsal view, the others are lateral views) and 41% (23/56) had a severe curvature (severe, the bottom one is a dorsal view). K) Image of a scoliotic *rpgrip1l*^Δ/+^ fish injected with the foxj1a::urp2 construct. Graph showing that among the survivor fish, 2/17 (12%) became scoliotic whereas none of the non-injected *rpgrip1l*^Δ/+^ heterozygotes became scoliotic.

## DISCUSSION

The study of juvenile scoliosis in many ciliary gene mutants is complicated by the occurrence of embryonic curvature and larval death, which requires rescuing embryonic defects to allow the mutants to reach adulthood [2]. In this paper we dissected the mechanisms of scoliosis appearance in zebrafish *rpgrip1l* mutants. *RPGRIP1L* is a causal gene in syndromic ciliopathies and its invalidation in humans and in mice leads to severe embryonic anomalies in several organs including the brain [5,11,12,28]. The absence of embryonic defects of *rpgrip1l*^Δ/Δ^ zebrafish, likely due to mRNA maternal stores [29] and/or to genetic compensation, makes it an excellent model to study scoliosis development.

Ventricular and CC dilations in *rpgrip1l*^Δ/Δ^ zebrafish mutants correlated with, or just preceded, the onset of scoliosis. These dilations could be the cause of *foxj1a* overexpression, since *foxj1a* is upregulated in response to epithelial stretch in the zebrafish pronephric tubules and to tissue injury in the spinal cord [30]. The defect in RF maintenance is also a likely consequence of cilia defects and ventricle dilations. In zebrafish embryos, the RF depends on cilia for its polymerization [3]. However, the mechanisms of RF maintenance have remained elusive, mainly due to the difficulty in visualizing this structure beyond embryonic stages. Our work allows visualizing the RF in juveniles and adults and demonstrates that SCO-spondin production and RF polymerization are continuous processes. In *rpgrip1l*^Δ/Δ^, the RF fails to be maintained at the onset of scoliosis. Given that SCO-spondin mutants develop spine deformities [31], the loss of the RF in *rpgrip1l*^Δ/Δ^ juveniles is a likely cause of scoliosis appearance.

The knock-down of *urp1* and *urp2* genes in zebrafish embryos leads to a curled down axis, while their overexpression leads to the opposite curvature [4]. *urp1* and *urp2* are expressed in a ventral population of spinal CSF-cNs which detect spinal curvature and modulate locomotion and posture by projecting onto spinal interneurons and motoneurons [32,33]. The upregulation of *urp1* and *upr2* expression in *rpgrip1l^Δ/Δ^* juveniles is consistent with the tail-up phenotype observed at scoliosis onset and suggests that both increased and reduced activity of this neuropeptide family in CSF-cNs can trigger scoliosis. Our finding that overexpressing *urp2* along the central canal is able to drive scoliosis in *rpgrip1l* heterozygotes confirms this hypothesis.

Our results pave the way for the analysis of the interactions between RF defects, URP signaling, *foxj1* overexpression and inflammation in scoliosis. Interestingly, neuroinflammation has been observed in *ptk7^-/-^* scoliotic fish and, in this model, scoliosis is alleviated by antiinflammatory treatments [20]. Moreover, inducing focal inflammation in zebrafish leads to scoliosis [20]. How inflammation is induced in *rpgrip1l* mutants requires further investigation. Cilia defects can trigger inflammation in the kidney and in the brain [34,35]. Moreover, Urotensin II signaling induces inflammation in many contexts [36], raising the interesting possibility that increased URP signaling from CSF-cNs could drive neuroinflammation in the *rpgrip1l*^Δ/Δ^ zebrafish scoliosis model.

In conclusion, our data support the idea that an upregulation of URP signaling in CSF-cNs, downstream of defects in CSF flow and RF maintenance, underlies scoliosis onset in *rpgrip1l* mutants. They allow us to propose that inappropriate levels of URP neuropeptides in these neurons, resulting in imbalanced regulation of posture or locomotion, may constitute an initial trigger of 3D spine deformities at a stage of intense body growth and tissue remodeling. This hypothesis is attractive given that loss of spine alignment is an intrinsic feature of scoliosis.

## ACKNOWLEDGEMENTS

We are grateful to the IBPS aquatic animal and imaging facilities and the ICM sequencing facility for their technical assistance. We thank Michaël Trichet from the IBPS electron microscopy facility for participating in the MEB experiments, the TACGENE facility for providing the Cas9 protein; the TEFOR Paris-Saclay facility for the brain clearing experiment; Thierry Jaffredo and Pierre Charbord for their precious help in transcriptome analysis; Nicolas Baylé for initial characterization of the *rpgrip1l* mutant; Claire Wyart for advice and critical reading of the manuscript. This work was supported by funding to SSM from the Fondation pour la Recherche Médicale (Equipe FRM EQU201903007943) and the Fondation Yves Cotrel.

## AUTHOR CONTRIBUTIONS

Conceptualization, C.V. and S.S.M.; Methodology, C.V., I.A., A.J., G.P. and C.P.; Investigation: C.V., I.A., G.P., Y.C.B., A.E., M.D., D.L.S., H.L.R. and J.V.; Formal analysis: C.V., H.K., A.J., and S.S.M.; Validation: C.V. and S.S.M.; Writing – Original draft, C.V. and S.S.M.; Funding acquisition, S.S.M.

## STAR* METHODS

### EXPERIMENTAL MODEL AND SUBJECT DETAILS

#### Zebrafish

Wild-type, *rpgrip1l^ex4^* and *rpgrip1l*^Δ^ zebrafish embryos and adults were raised, staged and maintained as previously described [37]. All our experiments were made in agreement with the european Directive 210/63/EU on the protection of animals used for scientific purposes, and the French application decree ‘Décret 2013-118’. The projects of our group have been approved by our local ethical committee ‘Comité d’éthique Charles Darwin’. The authorization number is 2015051912122771 v7 (APAFIS#957). The fish facility has been approved by the French ‘Service for animal protection and health’ with approval number A-75-05-25. All experiments were performed on Danio rerio embryos of mixed AB/TL background. Animals were raised at 28.5°C under a 14/10 light/dark cycle.

### CONTACT FOR REAGENT AND RESOURCE SHARING

Sylvie Schneider-Maunoury

Christine Vesque

### KEY RESOURCES TABLE

Provided separately

### METHOD DETAILS

#### Rpgrip1l mutant generation and genotyping

##### Guide RNA preparation and microinjection

Crispr target sites were selected for their high predicted specificity and efficiency using the CRISPOR online tool [38]. Real efficiency was assessed on zebrafish embryos by T7E1 test [39]. The two most efficient guides (Rpgrip1l_x4_G1: GCTTACGGTCCTTCACCAGACGG and Rpgrip1l_x25_G3: CCTCAGTTGACAGGTTTCAGCGG) respectively situated 24 nt from beginning of exon 4 and 82 nt downstream of exon 25 were kept for further experiments.

sgRNA transcription templates were obtained by PCR using T7_Rpgrip1l-x4_G1_Fw primer (5’-GAAATTAATACGACTCACTATAGGCTTACGGTCCTTCACCAGAGTTTTAGAGCTAGAAATAGC-3’) or T7_Rpgrip1l-x25_G3_Fw (5’-GAAATTAATACGACTCACTATAGGCCTCAGTTGACAGGTTTCAGGTTTTAGAGCTAGAAATAGC-3’) as forward primer and sgRNA_R universal primer (5’-AAAAGCACCGACTCGGTGCCACTTTTTCAAGTTGATAACGGACTAGCCTTATTTTAACTTGCTA TTTCTAGCTCTAAAAC-3’) as reverse primer. sgRNAs were transcribed using Megascript T7 Transcription Kit (Thermo Fisher Scientific, Waltham, MA) and purified using NucleoSpin^®^ RNA Clean up XS kit (Macherey Nagel, Düren, Germany). sgRNA:Cas9 RNP complex was obtain by incubating Cas9 protein (gift of TACGENE, Paris, France) (7.5 μM) with sgRNA (10 μM) in 20 mM Hepes-NaOH pH 7.5, 150 mM KCl for 10 min at 28 °C. 1–2 nl was injected per embryo. For deletion, Rpgrip1l_x4_G1 and Rpgrip1l_x25_G3 RNP complexes were mixed half and half.

##### Screening and genotyping

Injected (F0) fish were screened for germline transmission by crossing with wild type fish and extracting genomic DNA from obtained embryos. For genomic DNA extraction, caudal fin (juveniles/adults) or whole embryo DNA were used. Genomic DNA was isolated with Proteinase K (PK) digestion in 40 of lysis buffer (100 mM Tris-HCl pH 7.5, 1 mM EDTA, 250 mM NaCl, 0.2% SDS, 0. 1 μg/μl Proteinase K) for embryos (300 μl for adult fin) overnight at 37°C with agitation. PK enzyme was inactivated 10 min at 90°C and a five fold dilution was used as a template for PCR amplification. Genotyping of mutations in exon 4 and deletion between the two Crispr target sites was performed by PCR using Rpgrip1l_x4_F (TTGTGACACCGCATGCATTT) as forward primers and Rpgrip1 l_x4_R4 (CCGTGAGGTGGTCACCGG) as reverse primers for mutations and Rpgrip1 l-x25-R (ACATCAGAGGAAATCTTCTTTATTCAGC) as reverse primers for deletion.

#### μCT scans

The samples were scanned on a Bruker micro scanner (Skyscan 1272) with a resolution of 8.5 μm, a rotation step of 0.55° and a total rotation of 180°. For the acquisition of adult fish (2,5 cm), a 0.25 mm aluminium filter was used, for a voltage of 50 kV and an intensity of 180 mA, for juveniles fish (0,9 to 1,2 cm) the filter was omitted, and a voltage of 60 kV was used with an intensity of 166 mA. Each image contained 1008 x 672 pixels and was based on the average of 3 images. The 3D reconstruction by backprojection was carried out by the NRecon software and the 2D image overlays were then cleaned by the CT Analyser software. The Dataviewer software allowed all fish skeletons to be oriented in the same way taking the otoliths as landmarks. The CTvox software then allowed 3D visualizing of the samples. Morphometric analysis was performed with the CT Analyser software.

#### Scanning electron microscopy

Fish were euthanized using lethal concentration of MS222 (0.028 mg/mL). The brains were quickly dissected in 1.22 X PBS (pH 7.4), 0.1 M sodium cacodylate and fixed overnight with 2% glutaraldehyde in 0.61 X PBS (pH 7.4), 0.1 M sodium cacodylate at 4°C. They were sectioned along the dorsal midline with a razor blade to expose their ventricular surfaces. Both halves were washed four times in 1.22 X PBS and post-fixed for 15 minutes in 1.22 X PBS containing 1% OsO4. Fixed samples were washed four times in ultrapure water, dehydrated with a graded series of ethanol and critical point dried (CPD 300, Leica) at 79 bar and 38 °C with liquid CO2 as the transition fluid and then depressurized slowly (0,025 bar/s). They were then mounted on aluminum mounts with conductive silver cement. Sample surfaces were coated with a 5 nm platinum layer using a sputtering device (ACE 600, Leica). Samples were observed under high vacuum conditions using a Field Emission Scanning Electron Microscope (Gemini 500, Zeiss) operating at 5 kV, with a 20 μm objective aperture diameter and a working distance around 3 mm. Secondary electrons were collected with an in-lens detector. Scan speed and line compensation integrations were adjusted during observation.

#### Histological sections of juvenile and immunofluorescence on sections

Juvenile and adult zebrafish were euthanized using lethal concentration of MS222 (0.028 mg/mL). Pictures and size measurements were systematically taken before fixation. Fish were fixed in Zamboni fixative [35 ml PFA 4 %, 7.5 ml saturated picric acid (1.2 %), 7.5 ml 0.2M Phosphate Buffer (PB)] [40] overnight at 4°C under agitation. Fish were washed with Ethanol 70% and processed for dehydration by successive 1 h incubation in Ethanol (3 x 70% and 2 x 100%) and Butanol (2 x 100%) at room T° under agitation, then for paraffin inclusion. 14 μm sagittal paraffine sections were obtained using a Leica RM2125RT microtome. Sections were deparaffinized and antigen retrieval was performed by incubation for 7 min in boiling citrate buffer (10 mM, pH 6). Immunofluorescence staining was performed as described previously [41]. The following primary antibodies were used: anti-RF, anti-Urotensin II, anti-Acetylated Tubulin, anti-Glutamylated Tubulin, anti-Arl13b. Corresponding primary and secondary antibodies are described and referenced in the Key Resources Table. For URP intensity comparison, a 10μm Z-projection (Max intensity) was obtained for each field and graded pixel intensities were visualized using the FIRE tool in FIJI software. Three classes of intensities were defined: “weak” corresponds to blue color and partial cytoplasm staining, “medium” corresponds to pink color and uniform cytoplasmic staining, “strong” corresponds to yellow color and staining of both cytoplasm and axon. Ventral CSF-cN density corresponds to the ratio of URP-positive cells over the number of DAPI stained nuclei lining the CC ventrally.

#### Immunofluorescence on whole embryos

Embryos from 24 to 40 hpf were fixed 4 hr to overnight in 4% paraformaldehyde (PFA) at 4°C. Larvae at 48 and 72 hpf were fixed 2 hr in 4% PFA and 3% sucrose at 4°C, skin from the rostral trunk was partially removed and yolk was removed. Samples from 24 to 40 hpf embryos were blocked overnight in a solution containing 0.5% Triton, 1% DMSO, 10% normal goat serum and 2 mg/mL BSA. Samples from 48 to 72 hpf larvae were blocked in 0.7% Triton, 1% DMSO, 10% NGS and 2 mg/mL BSA. Primary antibodies were incubated one to two nights at 4°C in a buffer containing 0.5% Triton, 1 % DMSO, 1 % NGS and 1 mg/mL BSA. All secondary antibodies were from Molecular Probes, used at 1:400 in blocking buffer, and incubated 2.5 hr at room temperature. The following primary antibodies were used for in toto immunohistochemistry with anti-Reissner fiber and anti-Acetylated-tubulin antibodies. Corresponding primary and secondary antibodies are described and referenced in the Key Resources Table. Whole mount zebrafish embryos (dorsal or lateral mounting in Vectashield Mounting Medium) were imaged on an Leica SP5 confocal microscope equipped with a 63X immersion objective. Images were then processed using Fiji.

#### Whole-mount brain clearing

Brains were dissected from 4-5 wpf size-matched (0.9-1.2 mm length) zebrafish after *in toto* fixation with formaldehyde. Whole–mount tissue clearing was performed following the zPACT protocol [42]. In brief, brains were infused for 2 days in hydrogel monomer solution (4% acrylamide, 0.25% VA-044, 1% formaldehyde and 5% DMSO in 1X PBS) at 4°C. Polymerization was carried out for 2h30 at 37°C in a desiccation chamber filled with pure nitrogen. Brains were transferred into histology cassettes and incubated in clearing solution (8% SDS and 200 mM boric acid in dH2O) at 37°C with gentle agitation for 8 days. Cleared brains were washed in 1X PBS with 0.1% Tween-20 (PBT) for 3 days at room temperature and kept in 0.5% formaldehyde, 0.05% sodium azide in PBT at 4°C until further processing. Brains were subsequently placed for 1h in depigmentation pre-incubation solution (0.5X SSC, 0.1% Tween-20 in dH2O) at room temperature. The solution was replaced by depigmentation solution (0.5X SSC, Triton X-100 0.5%, formamide 0.05% and H2O2 0.03% in dH2O) for 45 minutes at room temperature. Depigmented brains were washed for 4h in PBT and post-fixed (2% formaldehyde and 2% DMSO in PBT) overnight at 4°C.

#### Cleared brain immunostaining

Whole-mount immunolabeling of cleared brains was performed as described in the zPACT protocol with slight modifications. Briefly, brains were washed in PBT for one day at room temperature and blocked for 10h in 10% normal goat serum, 10% DMSO, 5% PBS-glycine 1M and 0.5% Triton X-100 in PBT at room temperature. Brains were washed again in PBT for 1h prior to incubation with anti-ZO-1 antibody (ZO1-1A12, Thermofisher, 1:150) in staining solution (2% normal goat serum, 10% DMSO, 0.1% Tween-20, 0.1% Triton X-100 and 0.05% sodium azide in PBT) for 12 days at room temperature under gentle agitation. Primary antibody was renewed once after 6 days of incubation. Samples were washed three times in PBT and thereafter incubated with Alexa Fluor 488-conjugated secondary antibody (A11001, Invitrogen, 1:200) for 10 days in staining solution at room temperature under gentle agitation. Secondary antibody was renewed once after 5 days of incubation. Samples were washed three times in PBT prior to a counterstaining with DiIC18 (D282, Invitrogen, 1 μM) in staining solution for three days at room temperature with gentle agitation. Samples were washed in PBT for 3h before mounting procedure.

#### Mounting and confocal imaging

Samples were placed in 50% fructose-based high-refractive index solution (fbHRI, see [42]) /50% PBT for 1h, then in 100% fbHRI. Brains were mounted in agarose-coated (1% in standard embryo medium) 60 mm Petri dish with custom imprinted niches to help orientation. Niche-fitted brains were embedded in 1% phytagel and the Petri dish filled with 100% fbHRI. fbHRI was changed three times before imaging until its refractive index matched 1.457. Images were acquired with a Leica TCS SP8 laser scanning confocal microscope equipped with a Leica HC FLUOTAR L 25x/1.00 IMM motCorr objective. Brains were scanned at a resolution of 1.74×1.74×1.74 μm (xyz) and tiled into 45 to 70 individual image stacks, depending on brain dimensions, subsequently stitched, using LAS X software.

#### Volumetric analysis of the posterior ventricles

The volumes of the posterior ventricles were segmented, reconstructed and analyzed using Amira for Life & Biomedical Sciences software (Thermo Fisher Scientific). In essence the ventricles volumes were manually segmented in Amira’s segmentation editor and subsequently refined by local thresholding and simplification of the corresponding surfaces. Volumes, which were open to the environment were artificially closed with minimal surfaces by connecting the distal-most points of their surface to the contralaterally corresponding points using straight edges. Due to the biologic variability of the sample population the overall size of the specimens needed to be normalized to keep the measured volumes comparable. For this spatial normalization one of the specimens was randomly selected from the wildtype population as template (1664, grey) and the ‘Registration’ module in Amira was used to compute region-specific rigid registrations for the other specimens, allowing for isotropic scaling only [details in supp. mat.]. For excluding the influence of the ventricular volumes, the registration was computed on the basis of the independent reference stain (DiIC18) (details in supp. mat.). Region specific volume differences between the mutant and wildtype population were evaluated on seven subvolumes of the posterior ventricles (details in supp. mat.).

#### RNA extraction for transcriptome analysis and quantitative RT-PCR

Juvenile and adult zebrafish were euthanized using lethal concentration of MS222 (0.28 mg/mL). Their length was measured and their fin cut-off for genotyping. For transcriptomic analysis, brain and dorsal trunk from 1 month juvenile (0,9 to 1,0 cm) zebrafish were dissected in cold PBS with forceps and lysed in QIAzol (QIAGEN) after homogenization with plunger pistons and 1ml syringes. Samples were either stored in QIAzol at −80°C or immediately processed. Extracts containing RNA were loaded onto QIAGEN-mini columns, DNAse digested and purified in the miRNAeasy QIAGEN kit (Cat 217004) protocol. Samples were stored at −80°C until use. RNA concentration and size profile were obtained on the Tapestation of ICM platform. All preparations had a RIN above 9.2. For Q-PCR from 3 weeks juveniles, whole fish minus internal organs were lysed in Trizol (Life technologies) using the Manufacturer protocole.

#### Quantitative PCR

The cDNA from the isolated RNAs was obtained following GoScript Reverse Transcription System protocol (Promega), using 4 μg of total RNA for each sample. The 20 μL RT-reaction was diluted 4 fold and 4μl was used for each amplification performed in duplicates. The primers used for qPCR were the following: urp2 (F: AGAGGAAACAGCAATGGACG; R: TGTTGGTTTTCTTGGTTGACG) [Zhang X et al., 2018], rpgrip1l (F: CAGACACCTGCTGGAGTTACA; R: TCCTGACTCACATCAAACGCA), mob4 (F: CACCCGTTTCGTGATGAAGTACAA; R: GTTAAGCAGGATTTACAATGGAG) or lsm12b F GAGACTCCTCCTCCTCTAGCAT and R GATTGCATAGGCTTGGGACAAC. The Q-PCR were performed using the StepOne real-time PCR system and following its standard amplification protocol (Thermo Scientific). Relative gene expression levels were quantified using the comparative CT method (2-ΔΔCT method) on the basis of CT values for target genes and either mob4 or lsm12b as internal controls.

#### RNA-sequencing and analysis

RNA-seq libraries were constructed at ICM according to standard Illumina protocols and sequenced on an Illumina NovaSeq 6000 to obtain about 40 megabases of sequence in total, both strands. Paired-end reads were aligned against the GRCz11 genome using the STAR RNA-seq aligner with default parameters outputting FastQ files using Partekflow™. Gene-expression levels in terms of transcripts per million were estimated by HTseq-count, and differentially expressed gene (DEG) analysis was carried out using ANOVA. The genes with a fold-change value of 2 or more and an adjusted p value of less than 0.01 were defined as significant DEGs. The list of differentially expressed genes was then generated. Gene Ontology GO analysis was performed using DAVID (https://david.ncifcrf.gov/). Principle component Analysis PCA and Hierarchical clustering were performed using Partek genomics suite™.

#### Generation and injection of the foxj1a::upr2 construct

To generate the *foxj1a::urp2* construct, the *urp2* ORF was amplified from 24hpf zebrafish cDNA (URP2_BamHI_F: gcgcgcGGATCCgtatctgtagaatctgctttgctgc, URP2_XbaI_R: gcgcgcTCTAGAggcagagggtcagtcgtgttat) and cloned into pME-MCS (tol2kit, [45]). The final transgene was then obtained by LR recombination (using the Gateway vector construction kit (Life Technologies)) with the entry plasmids p5E-fox1a (containing *foxj1a* regulatory sequences [2]. pME-*urp2 and* p3E-polyA into pDestTol2CG2 the destination vector (tol2kit). The vector contains and an additional cassette consisting of EGFP expressed under the control of *cmlc2* promoter which allows visualizing GFP+ hearts from 1 dpf in injected embryos expressing the transgene. A volume of 1 nl of a mix containing the circular plasmid at 10-15 ng/μl together with Tol2 transposase mRNA at 20ng/μl was injected at the 1 cell-stage or at the 2-4 cell-stage embryos from *rpgrip1l*^Δ/+^ x wt outcrosses. The proportion of very malformed embryos at 24 hpf (reduced head, short and highly twisted body) was high when injected at the 1 cell-stage (77% death before 5 dpf, 37/48 embryos) and lower when injected at the 2-4 cell-stage (27% death before 5 dpf, n=77/106 embryos). At 5 dpf, injected larvae that had properly inserted the transgene were selected based on green fluorescence in the heart. Injected larvae and non-injected control siblings were raised to adulthood for genotyping and scoliosis scoring.

### QUANTIFICATION AND STATISTICAL ANALYSIS

For all experiments the number of samples analyzed is indicated in the text and/or visible in the figures (scatter plots). Because the number of samples was generally too low to achieve normality of distribution (except in Supplementary Figure 1), a non-parametric Mann-Whitney test was performed for all quantifications. For cilia length and density in Supplementary Figure 1, both parametric (unpaired T test) and non-parametric (Mann-Whitney test) were performed and gave similar results. Statistical analysis was performed using the Prism software. ****: P value < 0.0001; ***: P value < 0.001; **: P value < 0.01; *: P value < 0.05.

## SUPPLEMENTARY MATERIAL

**Supplementary Figure 1: Production and characterization of the *rpgrip1l^ex4^* and *rpgrip1l*^Δ^ mutant fish.g**

A) Graph showing the exon-intron structure of the *Danio rerio rpgrip1l* gene. Two alleles were produced by CRISPR/cas9-mediated genome engineering, *rpgrip1l^ex4^* with a nonsense mutation in exon 4 and *rpgrip1l*^Δ^ with a large deletion encompassing most of the protein coding sequence. Both alleles had identical phenotype (panel D and data not shown). Only *rpgrip1l*^Δ^ was further studied. B, C) Absence of muscle pioneer defects. In *rpgrip1l*^Δ/Δ^ as in control (ctl) 24 hpf larvae, a normal distribution of muscle pioneer cells expressing both Engrailed (green) and Prox1 (magenta) is observed, and a characteristic chevron shape of the somites (Prox1+, Engrailed-fast muscle cells) is present in *rpgrip1l^Δ/Δ^* embryos (C) as in controls (B), indicating normal Hh signaling. D) Absence of laterality defects and of kidney cysts in *rpgrip1l*^Δ/Δ^ and *rpgrip1l^ex/ex4^* embryos. E-G) Absence of retinal defects and normal photoreceptor cilia in 5 dpf *rpgrip1l*^Δ/Δ^ larvae. Retinal sections of control (E) and *rpgrip1l*^Δ/Δ^ (F) were immunostained with Zpr3 (red, to label the external segment of photoreceptors) and Acetylated Tubulin (Ac Tub, green) to label cilia. Insets in E and F focus on the photoreceptor cilia. G) Graph of photoreceptor cilia length. Cilia are slightly longer in *rpgrip1l*^Δ/Δ^ (n= 101 cilia) than in control (n=131 cilia) retinas. H-K) Cilia of olfactory sensory neurons (H, I) and of neuromast hair cells (J, K) appear similar in control (H, J) and *rpgrip1l^Δ/Δ^* (I, K) 5 dpf larvae. L) Graph of cilia density in the CC of juvenile control (12 ROIs of 2 animals) and *rpgrip1l^Δ/Δ^* (26 ROIs of 2 animals) fish. Density is the number of cilia per 100 μm CC length. Although the cilia density varies along the length of the CC, it is statistically lower in *rpgrip1l*^Δ/Δ^ juvenile fish CC than in control CC. M) Graph of cilia length (in μm) in the dorsal and ventral CC of control (n=135 cilia of 2 fish) and *rpgrip1l*^Δ/Δ^ (n=199 cilia of 2 fish) *rpgrip1l*^Δ/Δ^juvenile fish. Cilia are longer in *rpgrip1l*^Δ/Δ^, particularly in the dorsal CC. Scale bars: 20 μm in B, C; 50 μm in E, F; 25 μm in H, I and 10 μm in J, K.

**Supplementary Figure 2. Visualization of the RF in embryos and adults.**

A-F) Immunofluorescence on whole 30 hpf control (A, C, E) and *rpgrip1l*^Δ/Δ^ (B, D, F) embryos to visualize the RF (anti-RF antibody) and cilia (Acetylated Tubulin Antibody). Nuclei are stained with DAPI. The RF is present and of normal appearance in the central canal of *rpgrip1l*^Δ/Δ^ embryos. G-J) SEM on wt adult brain to visualize the RF in brain ventricles. Scale bars: 10 μm in A-F, 2 μm in G, 10 μm in H, 1 μm in I and 400 nm in J.

**Supplementary Movie 1. μCT in juveniles.**

Superposition of 3D-reconstructed spines of 5 wpf juvenile fish, 1 control (white), 3 *rpgrip1l*^Δ/Δ^ of different severities: 2 tail-up (yellow, blue) and 1 scoliotic (pink).

**Supplementary Movie 2. Ventricle reconstruction: whole brain view.**

**Supplementary Table 1. Excel file with a list of genes deregulated in *rpgrip1l*^Δ/Δ^.**

Sheet 1: Trunk transcriptome, list of genes upregulated in *rpgrip1l*^Δ/Δ^ samples compared to controls. Sheet 2: Trunk transcriptome, list of genes downregulated in *rpgrip1l*^Δ/Δ^ samples compared to controls. Sheet 3: Brain transcriptome, list of genes upregulated in *rpgrip1l*^Δ/Δ^ samples compared to controls. Sheet 4: Brain transcriptome, list of genes downregulated in *rpgrip1l*^Δ/Δ^ samples compared to controls. Data were filtered with p values < 0.01 for transcripts regarded as statistically significant and fold changes f ≥ 2. In column 5 (Refseq protein), Refseq is given for the human orthologue when available.

**Supplementary Table 2 List of Foxj1a target genes upregulated in the trunk of *rpgríp1l*^Δ/Δ^ juvenile fish.** The gene list was compared to Foxj1 target genes identified in [17–19]. Colour code: Yellow indicates that the target gene was found in [17] or [18] and in [19]; green indicates that the gene was found in [17] or [18] but not in [19]; orange indicates that it was found in [19] only.

**Supplementary Methods**

