## Supplementary material for "Loss of the Reissner Fiber and increased URP neuropeptide signaling underlie scoliosis in a zebrafish ciliopathy mutant": suppl. figures

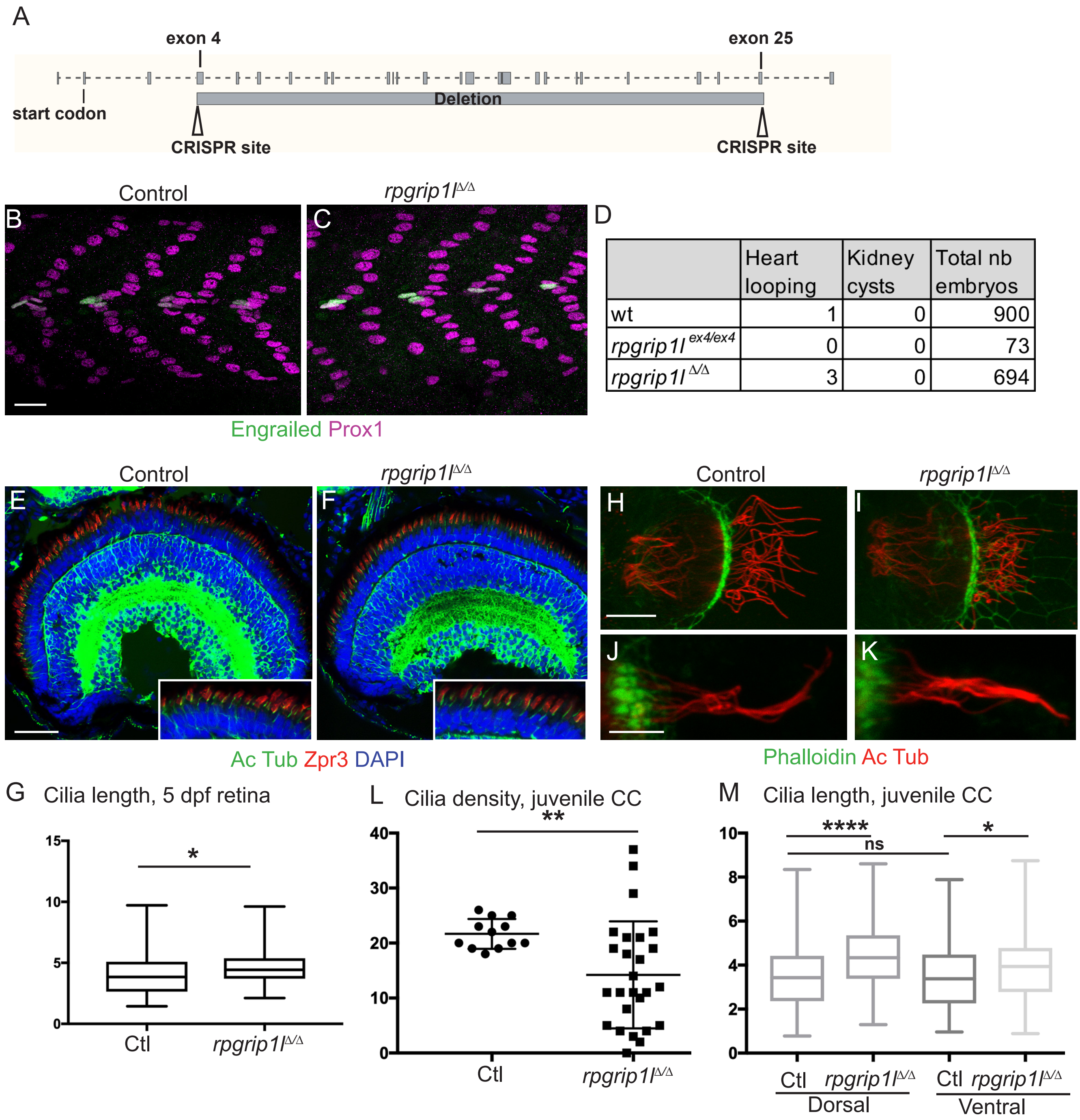


| | Heart looping | Kidney cysts | Total nb embryos |
|---|---|---|---|
| wt | 1 | 0 | 900 |
| *rpgrip1l* $^{ex4/ex4}$ | 0 | 0 | 73 |
| *rpgrip1l* $^{\Delta/\Delta}$ | 3 | 0 | 694 |

Vesque et al., Supplementary Figure 1

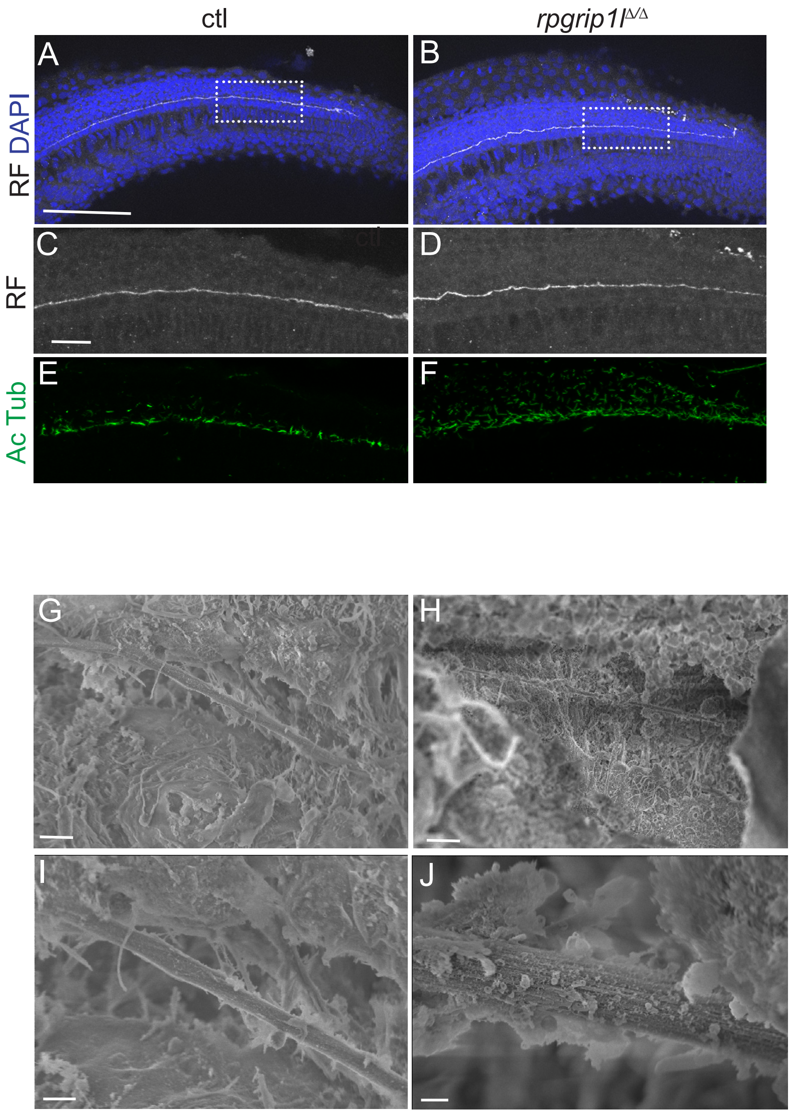


Vesque et al., Supplementary Figure 2
